# The Neural Architecture of Executive Functions Is Established by Middle Childhood

**DOI:** 10.1101/251447

**Authors:** Laura E. Engelhardt, K. Paige Harden, Elliot M. Tucker-Drob, Jessica A. Church

**Author notes:** Corresponding author: Laura E. Engelhardt, 108 E Dean Keeton St., Stop A8000, Austin, TX 78712.

## Abstract

Executive functions (EFs) are regulatory cognitive processes that support goal-directed thoughts and behaviors and that involve two primary networks of functional brain activity in adulthood. The current study assessed whether the same networks identified in adulthood underlie child EFs. Using task-based fMRI data from a diverse sample of *N* = 117 children and early adolescents (*M* age = 10.17 years), we assessed the extent to which neural activity was shared across three EF domains and whether these patterns reflected quantitative or qualitative differences relative to previously reported adult findings. Brain regions that were consistently engaged across switching, updating, and inhibition tasks closely corresponded to the cingulo-opercular and fronto-parietal networks identified in studies of adults. Isolating brain activity during more demanding task periods highlighted contributions of the dACC and anterior insular regions of the cingulo-opercular network. Results were independent of age and time-on-task effects. These results indicate that the two core brain networks that support EFs are in place by middle childhood. Improvement in EFs from middle childhood to adulthood, therefore, are likely due to quantitative changes in activity within these networks, rather than qualitative changes in the organization of the networks themselves. Improved knowledge of how the brain’s functional organization supports EF in childhood has critical implications for understanding the maturation of cognitive abilities.

## The Neural Architecture of Executive Functions Is Established by Middle Childhood

Cognitive maturation involves transitioning from stimulus-driven and reflexive actions to more deliberate thoughts and behaviors (1–3). Executive functions (EFs) – regulatory processes that monitor goal-directed cognitive operations – are critical for the developmental transition to adultlike thoughts and behaviors. Because of the importance of EFs for psychiatric health and cognitive skill formation in both childhood and adulthood (4–7), neuroscientists have been interested in understanding the neural mechanisms underlying normative maturation in EFs (8, 9). An exciting open question in this area is how the brain changes over development to support better performance across a variety of executive domains.

## Neural Architecture and Factor Structure of Executive Functions in Adulthood

Substantial individual differences and developmental differences are evident across separable EF domains, which include (a) *response inhibition*, or the ability to refrain from executing a practiced response; (b) *switching*, which requires performance adaptations in response to changing rules or goals; and (c) *updating*, which involves replacing information in working memory based on new demands (for reviews, see 10–12). Although these domains are statistically distinguishable, they also covary strongly, suggesting that domain-general executive resources underlie ability in any one specific domain. This pattern of relationships between EF domains is often referred to as the “unity and diversity” model (13).

Consistent with the “unity” of adult EFs, neuroimaging studies in adulthood have identified a core set of brain networks that are consistently activated in response to an array of tasks tapping different EF domains. Lesion studies and early functional magnetic resonance imaging (fMRI) work provided initial evidence that the prefrontal cortex (PFC) was fundamental to attention, working memory, and inhibition (for a review, see 14). More recent investigations employing multiple tasks have uncovered complex and distributed networks of brain regions active during EF tasks. Specifically, the *fronto-parietal network* includes bilateral inferior frontal gyrus (IFG), dorsolateral prefrontal cortex (dlPFC), inferior parietal lobule (IPL), superior parietal lobule (SPL), and pre-motor areas (14–19), and the *cingulo-opercular network* includes dorsal anterior cingulate (dACC) and bilateral anterior insula and is reliably active during error processing and task maintenance (20, 21).

Resting-state fMRI analyses suggest that findings from EF task-based studies identify networks of regions that are also intrinsically connected, as region-to-region correlations in spontaneous BOLD activity also cluster into dissociable fronto-parietal and cingulo-opercular networks across many samples (22–25). Overall, neuroimaging studies of adults have revealed a highly consistent set of regions that co-activate in response to executive demands. This detailed characterization sets a standard for evaluating the consistency of children’s EF-related brain activation.

## From Childhood to Adulthood: Qualitative or Quantitative Changes?

How do children’s brains mature to support improvements in EFs across development? Empirical results that answer this question will undoubtedly inform the design and evaluation of interventions to support children with EF deficits. One possible mechanism for age-related improvements in EF is that the networks of regions that support optimal deployment of executive control are not yet in place in childhood and that the maturation of EF results from the progressive establishment of an adultlike EF network over development. Support for such a *qualitative* account would come from findings that patterns of brain activity during executively demanding tasks are more diffuse among children or entirely distinct from patterns observed among adults. One example of qualitative, age-related changes in neural organization is early visually guided behaviors, which initially rely on subcortical activity before transitioning to predominantly posterior, and then anterior, cortical activation (26). Alternatively, a relatively consistent set of brain regions might undergo *quantitative* maturation before reaching their apex in adolescence or adulthood. This account of brain-behavior development would reflect strengthening or refinement of region-to-region connections and would be evidenced by engagement of a consistent set of brain regions across developmental stages (27). Declarative memory, for example, is mediated by activation in the medial temporal lobes and PFC from childhood through adulthood, with memory enhancement linked to age-related differences in the strength – but not location – of BOLD activity (28).

Behavioral studies of the factor structure of EF performance in childhood provide indirect support for quantitative maturation, i.e., that the neural architecture underlying successful engagement of executive resources is consistent across development. Notably, the “unity and diversity” model seen in adults, with a highly heritable factor that contributes to EF ability across domains and tasks, is evident as early as 8 years old (29). This suggests that common causal processes act on individual EFs in childhood, which is consistent with reliable, cross-task brain activity observed in adults.

Additionally, neuroimaging studies of EFs in childhood have found that individual tasks consistently engage temporal cortex, parietal cortex, and subcortical regions (e.g., 30–33). In conjunction with resting-state analyses (34), single-task studies highlight children’s engagement of the fronto-parietal and cingulo-opercular regions described above. However, a major limitation of neuroimaging studies of childhood EFs, compared to research in adults, is that they typically employ only a single EF task. Consequently, it is difficult to generalize findings across samples employing different tasks and to identify the extent to which task-related brain activity is task‐ or domain-specific versus general across EF domains.

To date, meta-analyses of children’s fMRI data have been the only avenue for addressing these questions. An early meta-analysis of 25 studies found evidence for consistent activation of bilateral prefrontal cortex, bilateral insula, and left parietal regions across tasks and age, as well as age-related changes in the lateralization of insula activity during EF individual tasks (35). More recently, a meta-analysis of 53 studies of individual EF tasks found evidence for cross-domain engagement of bilateral frontal, bilateral insula, and right parietal clusters, as well as evidence for domain-specific activation during switching and updating tasks (36). The regions identified in meta-analysis are largely consistent with the adult “core control system” described by Dosenbach and colleagues (20, 22), though with less consistency regarding the contribution of parietal regions.

However, meta-analyses cannot completely control for between-samples differences that may confound the results. For example, the greater number of studies examining the updating and inhibition domains, relative to the switching domain, may have biased previous findings regarding the relationships between these core constructs (36). Thus, meta-analyses of single-task studies provide circumstantial evidence suggesting that children activate a common set of brain regions during a variety of EF tasks and that these regions are the same as those activated by adults.

## Goals and Methodological Advantages of the Current Study

The goal of the current study was to provide the first direct test of whether the neural architecture of EFs in childhood is qualitatively the same as in adulthood. We hypothesized that the same functional brain networks that have been implicated in the adult literature (i.e., fronto-parietal and cingulo-opercular networks) would activate across three tasks tapping three distinct EF domains: switching, inhibition, and updating. To address this goal, we measured neural response to three EF tasks in a large, population-representative, and well characterized sample of children. This approach has several methodological advantages over previous meta-analytic approaches, including (a) the removal of between-study differences as a source of confounding variance; (b) the ability to apply greater quality control methods, including performance-based exclusionary criteria to isolate EF-related activity from noise; and (c) the ability to control for performance differences that may impact task-related fMRI signals. This is important because trial-by-trial variation in response time (RT), or “time-on-task” effects, positively correspond to activation in regions implicated in EFs, such as bilateral insula and right dlPFC (37). We address this issue by controlling for time-on-task effects at the whole-brain level and by separately examining the consistent BOLD correlates of RT across tasks. Finally, we are the first to conduct a formal comparison of activity in our sample to *a priori* regions from the adult literature.

## Results

### Task Performance

Means and standard deviations for task performance appear in Table 1. Table 2 reports correlations among these variables; as expected, performance covaried across tasks. Age was significantly associated with switching accuracy (*b* = .31, *SE* = .07, p < .001), updating hits-minus-false-alarms (*b* = .18, *SE* = .07, *p* < .05), and updating response time (*b* = -.18, *SE* = .07, *p* < .05). Age did not significantly predict switching response time (*b* = -.12, *SE* = .08, *p* = .13), inhibition accuracy (*b* = .12, *SE* = .08, *p* = .15), or inhibition response time (*b* = -.14, *SE* = .08, *p* = .09). Task performance differed by sex for updating response time, such that males responded .07s more quickly than females on average (*b* = -.45, *SE* = .20, *p* < .05). Inhibition response time also significantly differed by sex, such that females responded .02s more quickly than males on average (*b* = .41, *SE* = .20, *p* < .05).

**Table 1.** Descriptive statistics for task performance Domain: Performance measure

| Domain: Performance measure | *M* | *SD* |
| --- | --- | --- |
| Switching: Accuracy | .84 | .10 |
| Switching: Mean response time, correct trials | 1.11s | .15s |
| Updating: Hits minus false alarms | 7.71 | 3.95 |
| Updating: Mean response time, correct trials | .84s | .16 |
| Inhibition: Accuracy, go trials | .87 | .08 |
| Inhibition: Stop signal response time | .25s | .05 |

**Table 2.** Task performance correlations

|  | Switching<br>accuracy | Switching<br>RT | Updating<br>hits minus<br>false alarms | Updating<br>RT | Inhibition<br>accuracy | Inhibition<br>SSRT |
| --- | --- | --- | --- | --- | --- | --- |
| Switching<br>accuracy |  | -.48*** | .41*** | -.33*** | .44*** | -.11 |
| Switching RT | -.56*** |  | -.22* | .38*** | -.37*** | .01 |
| Updating hits<br>minus false alarms | .44*** | -.24* |  | -.38*** | .37*** | -.13 |
| Updating RT | -.41*** | .37*** | -.42*** |  | -.35*** | .08 |
| Inhibition<br>accuracy | .52*** | -.37*** | .41*** | -.41*** |  | -.02 |
| Inhibition SSRT | -.29*** | .24* | -.12 | .11 | -.13 |  |

*Note.* Zero-order Pearson correlation coefficients. Correlations using raw values are below the diagonal; those using age- and sex-residualized values are above the diagonal. *RT* = response time. See text for detailed description of measures.
\* $p \leq .05$ , \*\* $p \leq .01$ , \*\*\* $p \leq .001$

### Whole-brain Summed Mask Results

#### EF Contrasts

To examine the extent to which EF-related activity overlapped across tasks at the group level, we binarized the thresholded and cluster-corrected *z*-stat map for each task, then added the individual maps together to visualize areas of activation common across the three tasks (Figure 1). Significant task-positive activity common across all tasks was observed in the dorsal anterior cingulate cortex (dACC), bilateral anterior insula, right dorsolateral prefrontal cortex (dlPFC), bilateral inferior frontal gyri (IFG), bilateral frontal eye fields (FEF), bilateral superior parietal lobules (SPL), and bilateral anterior parietal cortex. Details on these regions can be found in Table 3. SI Appendix Figure S2a depicts areas of activity unique to each of the tasks, in addition to task-common regions. Significant task-negative activity (i.e., convergence of activity for baseline greater than EF-demanding conditions) is depicted in SI Appendix Figure S3.

**Figure 1.**
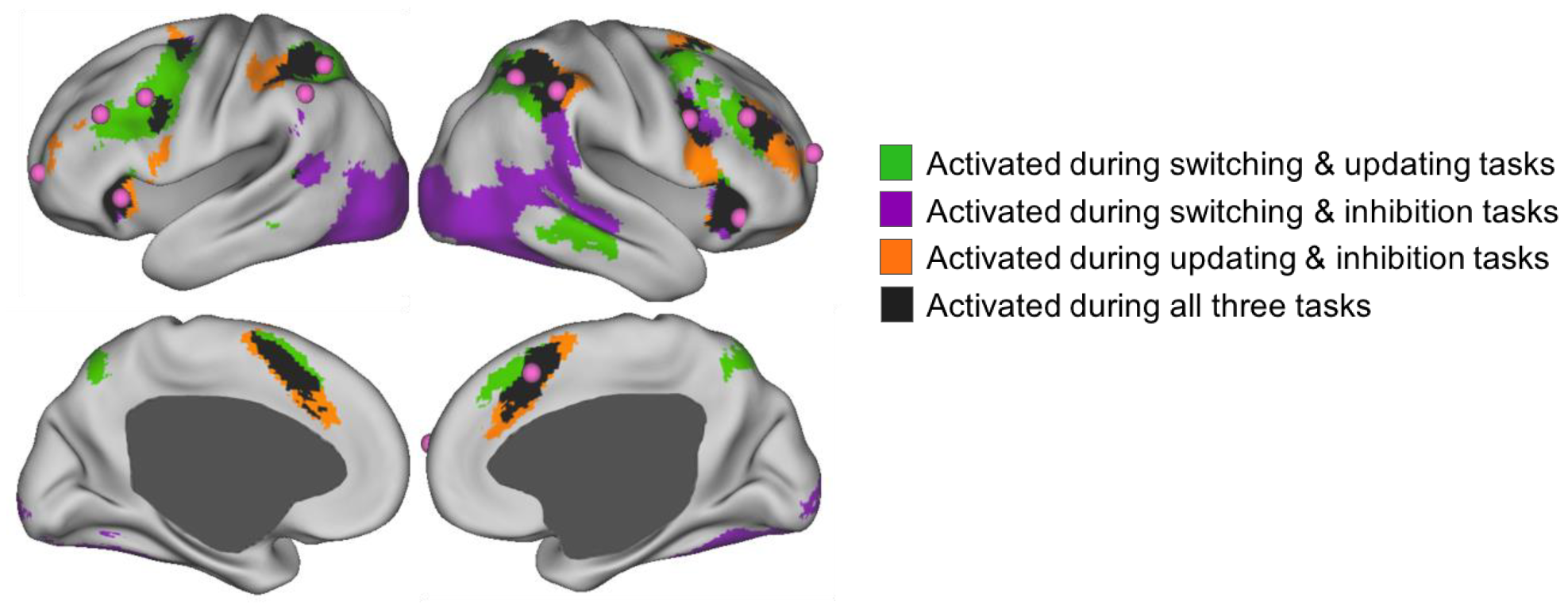
Overlapping task-positive brain activity across three EF tasks, overlaid with adult ROIs. Selected contrasts were *cue period during correct switch trials vs. baseline* for the switching task, *2-back blocks vs. baseline* for the updating task, and *correct stop trials vs. baseline* for the inhibition task. Prior to binarizing and summing across tasks, individual task maps were thresholded at *z* > 2.3 with a cluster probability of *p* < .001. Adult ROIs in pink were drawn from Dosenbach and colleagues (22, 73, 74).

**Table 3.** Task-overlapping centers of activity for EF contrasts

| Region | MNI coordinates |  |  | Cluster size<br>(voxels) |
| --- | --- | --- | --- | --- |
|  | x | y | z |  |
| Dorsal anterior cingulate cortex | 2 | 12 | 50 | 1162 |
| Left anterior insula | -30 | 22 | 2 | 456 |
| Right anterior insula | 36 | 20 | 2 | 734 |
| Right dorsolateral prefrontal cortex | 42 | 34 | 30 | 383 |
| Left inferior frontal gyrus | -42 | 2 | 28 | 88 |
| Right inferior frontal gyrus | 44 | 6 | 32 | 368 |
| Left frontal eye field | -26 | -6 | 50 | 248 |
| Right frontal eye field | 24 | 0 | 48 | 166 |
| Left anterior parietal cortex | -48 | -38 | 48 | 141 |
| Right anterior parietal cortex | 50 | -42 | 50 | 444 |
| Left posterior parietal cortex | -30 | -48 | 44 | 479 |
| Right posterior parietal cortex | 32 | -52 | 46 | 688 |
| Medial parietal cortex | 12 | -64 | 48 | 59 |

*Note.* Cluster size was determined by applying the *cluster* command in FSL to a mask of non-contiguous areas of activation with 50 voxels or more. Voxel size: 2×2×2mm.

After excluding eight participants diagnosed with developmental and/or learning disorders, the location and extent of activation were nearly identical to those of the whole sample (SI Appendix Figure S2b). We therefore proceeded with the entire sample for subsequent analyses.

#### ROI Geospatial Comparison

Our primary aim with respect to the adult literature-derived regions of interest (ROIs) was determining whether they converged with clusters exhibiting cross-task activity in our developmental sample. Figure 1 displays the literature-derived ROIs in pink, overlaid on activity common across two or more of the EF tasks. Ten of the 13 ROIs fell within areas of task-overlapping activity: bilateral anterior insula, dACC, bilateral IFG, bilateral IPS, right inferior parietal lobule (IPL), and right dlPFC. The right anterior prefrontal cortex (aPFC) from the adult ROIs overlapped partially with a task-common cluster, whereas the left aPFC, left dlPFC, and left IPL ROIs did not converge with cross-task activity in our sample. Of these, the left dlPFC adult ROI fell within an area of activity common across the switching and updating, but not inhibition, contrasts.

To directly compare the location of adult ROIs to children’s task-common activity, we computed the distance between the 13 a priori ROIs and the centers of 13 clusters of activity from our child sample (coordinates listed in Table 3). The majority of centroids representing overlapping activity in our child sample were within 15mm of the adult-based ROIs (see SI Appendix Table S2). Specifically, child activation in dACC, bilateral anterior insula, right dlPFC, bilateral IFG, right IPL, and right IPS lay 10mm or less from corresponding adult regions. Child activation in left IPS and left IPL were 10-16mm from corresponding adult regions. The three regions derived from the child data that were more distal from adult ROIs were bilateral FEF (each approximately 25mm from the adult IFG ROIs) and medial parietal cortex (22mm away from the right IPS adult ROI).

#### Response Time Contrasts

We next applied the summed-mask approach to the response time vs. baseline contrasts for the switching and inhibition tasks (Figure 2, Table 4). Large clusters of significant activity related to response time were observed in dACC, bilateral anterior insula, left premotor cortex, left FEF, right primary motor cortex, and right inferior occipital gyrus. Smaller clusters were identified in right premotor cortex, bilateral IFG, left anterior fusiform gyrus, and right IPS. To summarize, regions exhibiting the most extensive task-common activity as a function of response time matched the core task-control network comprised of dACC and anterior insula, with the addition of regions commonly associated with task execution and RT (37).

**Figure 2.**
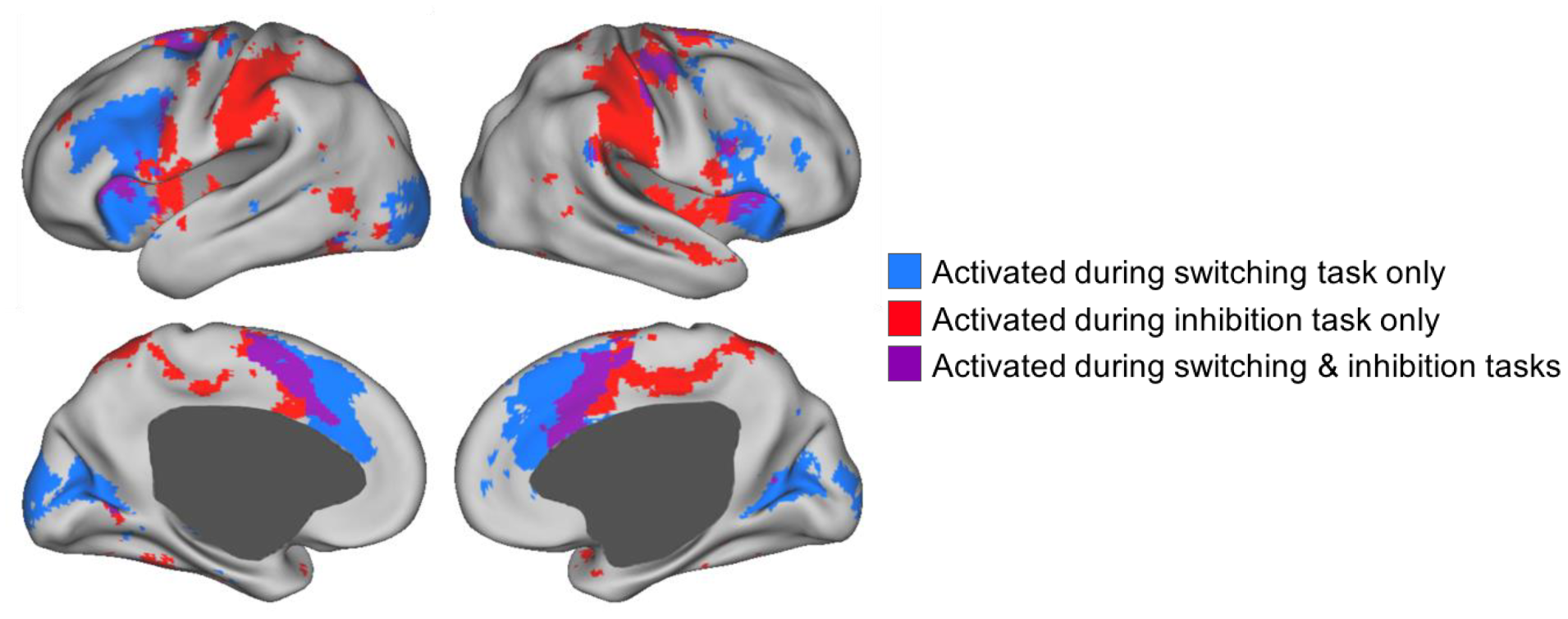
Overlapping brain activity corresponding to response time across two EF tasks. The contrast applied to both tasks was *mean-centered response time vs. baseline*. Prior to binarizing and summing across tasks, individual task maps were thresholded at *z* > 2.3 with a cluster probability of *p* < .001.

**Table 4.** Data-driven centers of activity for response time contrasts

| Region | MNI coordinates |  |  | Cluster size<br>(voxels) |
| --- | --- | --- | --- | --- |
|  | x | y | z |  |
| Dorsal anterior cingulate cortex | 0 | 4 | 50 | 1224 |
| Left anterior insula | -32 | 16 | 10 | 213 |
| Right anterior insula | 36 | 14 | 8 | 281 |
| Left inferior frontal gyrus | -52 | 8 | 24 | 124 |
| Right inferior frontal gyrus | 44 | 6 | 26 | 37 |
| Left premotor cortex | -20 | 0 | 62 | 93 |
| Right premotor cortex | 14 | -2 | 64 | 129 |
| Left frontal eye field | -26 | -8 | 54 | 141 |
| Right primary motor cortex | 38 | -16 | 56 | 289 |
| Right inferior occipital gyrus | 26 | -90 | -6 | 120 |
| Left intraparietal sulcus | -18 | -68 | 44 | 40 |

*Note.* Voxel size: 2×2×2mm.

### Sensitivity Analyses

#### Stricter Contrasts

We conducted a summed-mask analysis using the following more restrictive contrasts for the switching, updating, and inhibition domains, respectively: correct switch trials vs. correct repeat trials during the cue period, 2-back block vs. 1-back block, stop trials vs. correct go trials. As displayed in SI Appendix Figure S4, significant activity shared across all tasks was observed in small clusters within dACC (MNI coordinates: 0, 22, 44), right anterior insula (MNI coordinates: 32, 18, 8), right FEF (MNI coordinates: 22, 8, 48; 36 voxels), left IFG (MNI coordinates: -32, 8, 28), and left IPS (MNI coordinates: -46, -40, 52). The center of dACC activation was 7mm from the corresponding adult dACC region. Activation in the right insula, left IFG, and left IPS were 11, 13, and 18mm, respectively, from the corresponding adult ROIs. Right FEF activity was greater than 20mm from any adult region.

#### Age

To determine whether subtle age differences within the sample accounted for overlapping activation across EF tasks, we included mean-centered age as an independent variable in the GLM for each task, then applied the summed-mask approach to identify significant age-related activity common across EF domains. There were no significant clusters of age-correlated activity shared by the three tasks (Figure 3).

**Figure 3.**
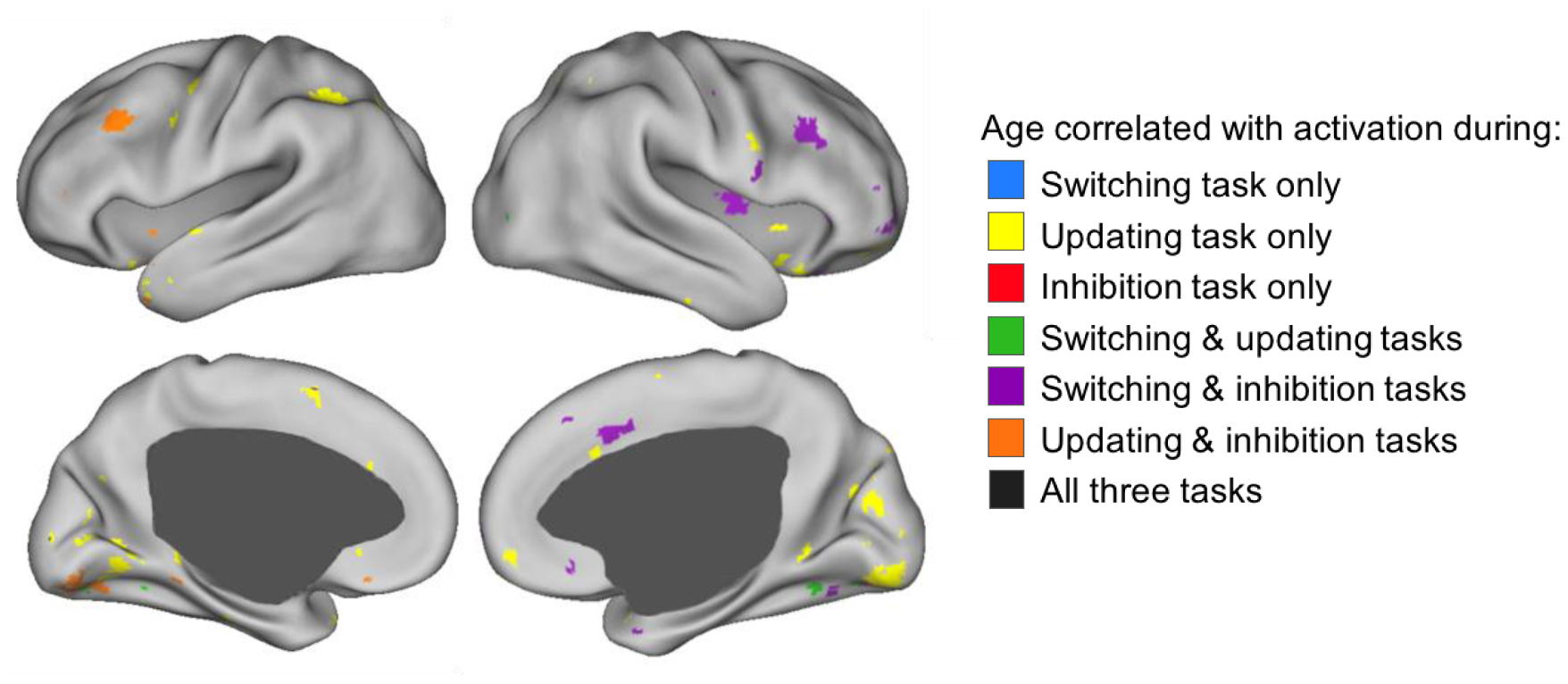
Overlapping task-positive brain activity as a function of age. Correlations between mean-centered age and percent signal change for the following contrasts: *cue period during correct switch trials vs. baseline* for the switching task, *2-back blocks vs. baseline* for the updating task, and *correct stop trials vs. baseline* for the inhibition task. Prior to binarizing and summing across tasks, individual task maps were thresholded at *z* > 2.3 with a cluster probability of *p* < .001.

## Discussion

Executive functions are foundational processes that underlie the development of complex reasoning and academic achievement (e.g., 4, 38–40). In addition to their impact on cognitive outcomes throughout development, EFs forecast psychological and physical wellbeing, and they mediate environmental risk for negative outcomes (4–7, 41). Understanding the neurobiological organization of EFs as they undergo rapid maturation in childhood is key to developing interventions that promote EF development, ameliorate executive deficits, and identify risk factors for impending cognitive and psychiatric impairments. An outstanding question is whether the functional brain networks that support domain-general EFs in adulthood are in place by middle childhood or whether they are substantively different. Motivated by well documented findings of task-overlapping activity in the adult literature (20, 23), meta-analyses of single-task studies of children’s EF-related activation (35, 36), and behavioral studies showing consistency in the factor structure of EF performance across development, we examined brain activation across switching, updating, and response inhibition tasks in a single, population-representative sample of children.

### Activity Shared Across EFs in Childhood Maps onto Adult Regions

Consistent with our hypotheses, we found that children engage a common set of brain regions across EF domains and that these regions overlap substantially with two EF networks that have been well characterized in adults. Specifically, regions that comprise the adult cingulo-opercular network (dACC, bilateral anterior insula) and the fronto-parietal network (right dlPFC, bilateral IFG, bilateral anterior and posterior parietal regions) consistently co-activated in response to EF demands in our child sample. Additionally, bilateral frontal eye fields at the intersection of the middle frontal and precentral gyri exhibited significant activation across all three tasks. Although FEFs are absent from task-related EF networks in studies of adults, these regions are functionally co-activated with regions in the adult fronto-parietal network during resting state (24, 25, 42–44). Our findings indicate that established patterns of neural activity underlying adult EFs are essentially in place by middle childhood. Thus, the development of EFs from middle childhood to adulthood likely involves quantitative changes in activity within EF-related networks, rather than qualitative changes in the organization of the networks themselves.

Employing stricter contrasts for the tasks revealed a specialized set of regions that were engaged across more demanding task periods. The largest clusters were centered upon dACC and right anterior insula, with less extensive cross-task activity observed in right FEF, left IFG, and left IPS. In adults, the dACC and anterior insula have been found to constitute a “core task-set system” based on their involvement across distinct trial periods and executive domains (20, 21). These results support the proposition that, in both childhood and adulthood, co-activation of cingulo-opercular regions is necessary for the execution of highly demanding tasks. Overall, the results of both the primary and sensitivity analyses imply that well documented increases in EF abilities from middle childhood to adulthood operate via a functional architecture that is in place by middle childhood.

Importantly, our key findings were independent of commonalities driven by response time, as trial-by-trial RT was included as a covariate in first-level analyses. This was a critical advantage of the current study, as meta-analyses of the neural basis of EFs in childhood are unable to dissociate activation attributable to executive processes per se from activation attributable to response time. The importance of this step was underscored by our finding that, across tasks, within-person differences in RT corresponded to activity in multiple regions relevant to EFs: dACC, bilateral anterior insula, bilateral IFG, left FEF, and left IPS. These patterns were highly consistent with RT effects observed in adult samples (37, 45). In summary, we found that activity in regions identified as critical to EF consistently relates to a standard behavioral outcome, but that activity in these regions also occurs above and beyond performance differences across individuals.

Evidence for commonalities in children’s brain activation across switching, updating, and inhibition tasks agrees with our and others’ findings that individual differences in EF task performance are best captured by a hierarchical model in which variance is shared across domain-specific EF factors, suggesting that common causal processes act on individual EFs (13, 29). Indeed, variance shared across EF domains is attributable primarily to genetic factors; a general factor of EF has been found to be nearly 100% heritable in samples of 7- to 14-year-olds (29) and young adults (46), with negligible contributions from environmental sources. Moreover, general intelligence, which overlaps strongly with a general factor of EF both phenotypically and genetically (47), exhibits high rank-order stability by age 10; predominantly genetic factors account for increasing stability in intelligence (48). Executive function thus constitutes one of the most stable and genetically influenced phenotypes early in life, and our finding that the neural architecture of EFs is effectively in place in middle childhood is consistent with behavior genetic evidence of maximum heritability in a general factor of EF at the same age.

### Implications for Future Research

The convergence of our results across brain activity and behavioral models clearly suggests that the organizational foundation for EFs is in place by middle childhood. The question that logically follows is: What mechanisms underlie the large gains in executive skills from middle childhood forward? One possibility is that changes in EF performance result from functional changes in the task-common regions we have described. For example, strength of co-activation between pairs or subsets of regions may increase with repeated engagement in EF-demanding situations over development (49). In the current study, we focused on global patterns of activation rather than inter-regional relatedness, but examining finer-tuned synchronicity between regions that exhibited significant activation across our tasks will likely prove fruitful. Another possibility is that structural maturation of brain regions and their connections mediates behavioral improvement in EFs. Exploring this possibility, Baum and colleagues (50) examined age-related changes in white matter-based connectivity and EF abilities in a cross-sectional sample of children through young adults. The degree to which white matter connectivity was stronger within functional modules (e.g., somatosensory regions, fronto-parietal regions) mediated developmental increases in performance on an EF task. The integration of functional and structural neuroimaging approaches would shed light on the mechanisms by which various neural properties interact to support the development of EFs.

Other extensions of this work may focus on the role of cingulo-opercular and fronto-parietal regions in the onset and maintenance of atypical thoughts and behaviors, as EF deficits have been implicated in nearly every developmental disorder (5, 51). The current results could provide a baseline against which studies of atypical development may be compared. For example, the regions highlighted in the current paper (in particular, dACC and bilateral anterior insula) show robust activation in the face of different executive demands. Hypo‐ or hyperactivation of these regions may therefore correspond to poor EF performance, as well as symptom burden. A recent meta-analysis of adult neuroimaging studies examined brain activity in response to EF tasks, comparing healthy controls to participants with various psychiatric disorders (52). Regardless of disorder type, EF-related activity among diagnosed groups consistently differed from that of healthy controls in left anterior insula, right vlPFC, right IPS, right motor regions, and anterior dACC. The authors proposed that brain networks that support adaptive cognitive control, like the fronto-parietal network, are especially vulnerable to disruptions that may manifest as psychopathology. Alternatively, divergence from established EF-related regions may be symptomatic of psychiatric or developmental disorders (53).

The current results tell us about developmental norms with respect to children’s brain organization. Future research that looks beyond group means may lead to greater understanding of the practical consequences and correlates of individual differences in engaging regions common across or unique to EF domains. For example, it may be that the regions we have identified here are required for successfully engaging in a task, whereas task-unique activation exhibits greater variability that may meaningfully relate to differences in task performance or other behavioral outcomes (54, 55).

### Limitations

We acknowledge a number of limitations in the current study, including a lack of collection of the same set of tasks in adults. However, adult EF activity has been well established across multiple tasks within large samples (e.g., 20, 23). By capitalizing on extant adult datasets, we were able to estimate the spatial proximity of hubs of activity in our sample to that of well characterized adult ROIs. The idiosyncrasies inherent to the tasks we selected constitute another limitation. In particular, the inhibition task led to strongly right-lateralized activation, potentially explaining the fewer left hemisphere overlaps across all three tasks. However, our tasks benefited from strong performance in the current sample, which is critical when interpreting developmentally normative brain activation during tasks, as error-related BOLD responses can differ systematically from more task-relevant signals (56, 57).

### Conclusion

The goal of this study was to evaluate the consistency of children’s brain activation in response to various EF demands and to determine whether co-activated regions followed the organization observed among adults. The study benefited from a large, representative sample measured on multiple tasks, conferring greater precision than that afforded by meta-analyses. The results indicated that, by middle childhood, a common set of fronto-parietal and cingulo-opercular regions support executive processing across EF domains. The results shed light on the neurobiological bases of a set of abilities that are critical for everyday functioning and lifelong wellbeing, indicating the organization is established by middle childhood. Further exploration of correlates of task overlapping and task unique EF-related signals presents an exciting opportunity to understand cognitive maturation in typical and atypical development.

## Materials and Methods

### Participants

As part of the neuroimaging arm of the Texas Twin Project (58), 127 twins or multiples in 3^rd^ through 8^th^ grade participated in an MRI session. Ten participants were excluded from the analyses due to incidental findings, equipment malfunction, refusal to continue, or failure to meet movement and performance cutoffs across all collected tasks. The final sample consisted of 117 participants with mean age of 10.17 years (*SD* = 1.37, range = 7.96 to 13.85); 57 participants were female. Participants reported diverse racial and ethnic backgrounds: 43.6% were non-Hispanic white, 14.5% were Hispanic, 5.1% were African American, 5.1% were Asian, 1.7% were another race, and 29.9% reported multiple races or ethnicities. The sample comprised 52 twin pairs (21 monozygotic, 16 same-sex dizygotic, and 15 opposite-sex dizygotic) and 13 individuals whose co-twins were not scanned. Zygosity was determined by a latent class analysis of researchers’ and parents’ ratings of twins’ physical similarity. The current study does not examine twin relations.

Developmental or learning disorder diagnoses were reported by parents for eight participants. Six participants had attention deficit hyperactivity disorder (ADHD), one of whom also reported non-specific reading disability; two had Asperger syndrome; and one had dyslexia. Results of the primary analyses with and without these individuals are described below.

### MRI Data Acquisition

Twins were scanned consecutively on the same day. Parents provided informed consent for their children’s participation, and participants provided informed assent. Participants were compensated for their time. Images were acquired on a Siemens Skyra 3-Tesla scanner with a 32-channel head matrix coil. We collected T1-weighted structural images with an MPRAGE sequence (TR = 2530 ms, TE = 3.37 ms, FOV = 256, 1×1×1mm voxels), as well as T2-weighted structural images with a turbo spin echo sequence (TR = 3200 ms, TE = 412 ms, FOV = 250, 1×1×1mm voxels). During tasks, we collected functional images using a multi-band echo-planar sequence (TR = 2000 ms, TE = 30 ms, flip angle = 60°, multiband factor = 2, 48 axial slices, 2×2×2mm voxels, base resolution = 128×128). Tasks were run on PsychoPy version 1.8 (59); stimuli were projected at a resolution of 1920×1080 to a screen that participants viewed via a mirror attached to the head coil. Participants wore Optoacoustics headphones and provided responses using a two-button response pad.

### fMRI Tasks

Task order was fixed to maximize the likelihood of retaining usable data across EF domains, and to avoid confounding sequence effects with individual differences (60). Tasks were ordered as follows: resting state (not presented here), switching task, updating task, inhibition task, switching task, updating task, resting state.

#### Switching task

Participants performed up to two runs of a cued switching task (SI Appendix Figure S1a; 61). Runs consisted of 46 trials in which participants were cued to pay attention to the shape or color of a target stimulus that would appear later. The two possible rules (shape and color) and two responses choices were displayed for the duration of the trial. A red box indicating which rule to follow appeared for the first 1.5 seconds of the trial. On 37 of the 46 trials, the target stimulus appeared .5 seconds after the red box disappeared, and the target remained on the screen for 2 seconds, during which time the participant could indicate which of the response choices matched the target. The response period was followed by a 1 second fixation cross. In 9 trials interspersed throughout the run, a target did not appear and a red fixation cross was displayed for .5 seconds, followed by a white fixation cross for .5 seconds. The cue-only trials allowed us to separate neural signals during the cue period from those during the target stimulus period (62). All trials were followed by a jitter of 0-8 seconds. The total run time was 5 minutes and 22 seconds. In the first run, the cued rule was consistent with the previous rule on 22 trials (repeat trial), and these were interspersed with 23 trials where the cued rule switched (switch trial). In the second run, there were 23 repeat rule trials, and 22 switch rule trials.

#### Updating task

Participants completed up to two runs of an N-back task (SI Appendix Figure S1b; adapted from 63). Each run consisted of 64 shape stimuli evenly divided into a 1-back and 2-back block. Block order was fixed. Prior to each block, participants viewed an instruction picture for 4 seconds that indicated whether they should look for shapes that matched one shape prior (1-back) or two shapes prior (2-back). During the blocks, each stimulus appeared for 1.5 seconds, followed by a 1 second inter-stimulus interval. A 20 second fixation followed each block. Each block had a total of 7 matches (21.9% of trials). Updating runs lasted 3 minutes and 32 seconds.

#### Inhibition task

To assess response inhibition, we administered one run of a visual Stop Signal task (SI Appendix Figure S1c; 64). Runs consisted of 96 “go” trials in which participants were instructed to indicate whether a horizontal arrow pointed to the left or the right, interspersed with 32 “stop” trials (25% of total trials) in which a red X appeared on top of the arrow, cueing the participant to withhold a respond. Across all trials, arrows were displayed for 1 second, with a 1 second interval, followed by a jittered fixation of 0 to 4seconds. For the first stop trial of each run, the X appeared .25 seconds after the arrow and remained on the screen for the duration of the arrow stimulus. If the participant correctly stopped on a given stop trial, the time between the appearance of the arrow and X on the next stop trial increased by .05 seconds; if the participant failed to inhibit a response, the time between the appearance of the arrow and X on the next stop trial decreased by .05 seconds. The inhibition task run lasted 6 minutes.

### Analyses

#### Behavioral analyses

To evaluate task performance, we selected one accuracy measure and one response time (RT) measure for each task. Variables of interest for the switching task were proportion of correct trials and mean RT for correct trials. Performance measures for the updating task were mean RT for correct trials and hits minus false alarms, or the difference between correct identification of *N*-back matches and misidentification of non-matches. Performance was collapsed across 1- and 2-back blocks. For the inhibition task, performance was evaluated by proportion of correct go trials and stop signal RT (SSRT), which estimates the time it takes to detect and correctly respond to (i.e., by inhibiting a response) a stop cue. The SSRT is determined by subtracting the mean time between presentation of the arrow and the red X from the mean RT for go trials.

Runs were excluded if performance did not meet the following criteria: for the switching task, at least 60% accuracy; for the updating task, at least four correct matches on 1-back blocks, 2 correct matches on 2-back blocks, and no more than 9 false alarms (indicating a match when there is none); for the inhibition task, selecting the correct arrow direction on 70% of trials or more, selecting the wrong direction on fewer than 10% of trials, stop accuracy between 25% and 75%, and stop signal reaction time greater than 50ms (65). Seventy-two runs (12.9% of total collected) were omitted for poor performance. Performance data were averaged across usable runs.

Analyses that included behavioral or demographic data were conducted in *R* version 3.2.3 (66). Statistical tests were conducted on standardized values. To account for the nonindependence of data drawn from individuals nested within families, we used the *nlme* R package to run regressions as linear mixed models with random intercepts.

#### fMRI preprocessing

Imaging data were preprocessed with the fMRI Expert Analysis Tool in FMRIB Software Library (FSL) version 5.9 (http://www.fmrib.ox.ac.uk/fsl). High-resolution T_1_-weighted structural images underwent skull stripping and brain extraction using Freesurfer version 5.3.0 (67). Functional data were registered to the structural image with a boundary-based algorithm (68), and structural images were registered to MNI space with the FMRIB Linear Image Registration Tool (69). Additional pre-statistics processing included spatial smoothing using a Gaussian kernel of FWHM 5mm; grand-mean intensity normalization of the 4D dataset by a single multiplicative factor; and high pass temporal filtering (Gaussian-weighted least-squares straight line fitting, with 50s sigma).

#### fMRI analyses

First-level analyses for on individual task runs were conducted with the FSL’s Improved Linear Model, which extends the voxelwise general linear model by estimating and correcting for time series autocorrelation (70). Data were modeled with a double-gamma HRF convolution. The highpass filter was set at 100s for the switching and inhibition runs and to 200s for the updating runs, the latter representing twice the duration of stimuli presentation. First-level models included six motion regressors; temporal derivatives for each regressor (except for the updating task, due to its block design); a trial-level response time regressor; and nuisance regressors that censored individual volumes identified to have excessive motion, defined as framewise displacement greater than .9mm (71). Two runs (.3% of total collected) were excluded from further analysis due to excessive motion during 60% of frames or more. Of the remaining usable runs, 11.0% of volumes were censored due to movement exceeding .9mm. Thirteen additional runs (2.3% of total collected) did not pass visual inspection at the registration stage and were omitted from subsequent analyses. In total, we retained 195 usable runs across 110 participants for the switching task, 170 usable runs across 100 participants for the updating task, and 100 usable runs across 100 participants for the inhibition task.

We selected contrasts that we anticipated would capture robust control-related activation for each task. The contrast for the switching task was the cue period during correct switch trials (i.e., when participants were cued to focus on a rule that differed from the previous trial) vs. baseline. For the updating task, the contrast was 2-back blocks vs. baseline. For the inhibition task, the selected contrast was correct stop trials vs. baseline. We also modeled response time vs. baseline contrasts for the switching and inhibition tasks to compare time on task effects to our principal results. The updating task could not be incorporated into the response time contrasts due to the nature of the block design.

Second-level analyses, which average contrast estimates over runs for each participant, were carried out by specifying a fixed effects structure within FMRIB Local Analysis of Mixed Effects (FLAME, 72). Third-level group analyses for each task were also executed using FLAME. To correct for whole-brain multiple comparisons, *z*-statistics were thresholded with a cluster-forming threshold of *z* > 2.3 (corresponding to voxelwise threshold of *p* < .01) and a cluster probability of *p* < .001 using Gaussian random field theory. We applied these relatively conservative cluster options to account for the increased likelihood of identifying false positives at the voxel and cluster level that arises from the nesting of multiple individuals within the same family.

#### Summed Masks Analysis

We first aimed to test the extent to which patterns of EF-related activation at the whole-brain level overlapped across tasks. Within the thresholded and corrected *z*-stat map for each task, we assigned a value of 1 to voxels that exhibited significantly (*z* > 2.3, *p* < .001) greater BOLD activity for the executive condition relative to baseline; voxels that failed to meet this criterion were assigned a 0. We added the binarized maps for each task together, resulting in a single map displaying voxels engaged by only one task, across two tasks, and across all three tasks. The results of this approach were compared to *a priori* regions of interest (ROIs) derived from the adult literature, as well as data-driven ROIs derived from the group results. For the response-time contrasts, thresholded and cluster-corrected maps for the switching and inhibition tasks were binarized and summed as described above.

#### ROI Geospatial Comparison

Clusters of activation derived from the summed mask analysis were compared to the location of 13 adult ROIs based on previous work examining domain-general task-control activity (22, 73, 74). These ROIs, listed along with their coordinates in SI Appendix Table S1, included five regions from the cingulo-opercular network and eight regions from the fronto-parietal network. In order to estimate distances between the literature-derived ROIs and the clusters of task-overlapping activity in the child sample, we identified the center of each cluster based on visual inspection of the summed mask (coordinates and cluster size provided in Table 3). For the current analyses, we considered only cortical clusters. The distance between each literature-derived and data-driven ROI was computed as:

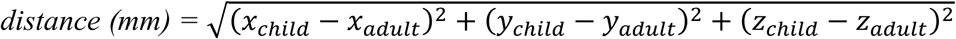

where x, y, and z correspond to the MNI coordinates for the child centers of activity and adult ROIs.

#### Sensitivity Analyses

We aimed to evaluate the generalizability of our results by rerunning key analyses on a more restricted set of contrasts. For the switching task, the selected contrast was correct switch trials vs. correct repeat trials during the cue period. For the updating task, the contrast was 2-back blocks vs. 1-back blocks. For the inhibition task, the selected contrast was correct stop trials vs. correct go trials. Because of the more constrained nature of these contrasts, the cluster-correction threshold was raised to *p* < .01.

To address the possibility that overlapping activity across tasks could be driven by within-sample age differences, we included mean-centered age as an independent variable in a final set of analyses, with the original contrasts and cluster threshold (*p* < .001) applied. The resulting binarized masks were then summed, revealing areas of the brain in which age significantly correlated with EF-related activation across the three tasks.

## Acknowledgements

This project was supported by National Institutes of Health grants R21 HD081437 (ETD and JAC) and R01 HD083613 (ETD). L. E. Engelhardt was supported by a National Science Foundation Graduate Research Fellowship. K. P. Harden and E. M. Tucker-Drob are each supported by Jacobs Foundation Research Fellowships. The Population Research Center at the University of Texas at Austin is supported by National Institutes grant R24 HD042849. We wish to thank the research team that facilitated data collection: Mary Abbe Roe, Jessica Graves, Annie Zheng, Damion Demeter, Saloni Kumar, Mackenzie Mitchell, and Tehila Nugiel. Finally, we thank participating families for their time and effort.

